# A single nonsynonymous mutation on gene encoding E protein of Zika virus leads to increased neurovirulence *in vivo*

**DOI:** 10.1101/2020.01.20.912535

**Authors:** Zhihua Liu, Yawei Zhang, Mengli Cheng, Ningning Ge, Jiayi Shu, Zhiheng Xu, Yigang Tong, Chengfeng Qin, Xia Jin

## Abstract

Zika virus can infect a wide range of tissues including the developmental brain of human fetuses, causing from mild to severe clinical diseases. Whether its genetic characteristics impacts on viral pathogenesis is incompletely understood. We have obtained viral variants through serially passage of a clinical Zika virus isolate (SW01) in neonatal mice *in vivo* and found some of which exhibited markedly increased virulence and neurotropism. By deep sequencing analysis, the more pathogenic viral variants were found to contain four dominant nonsynonymous nucleotide mutations on genes encoding E and NS2A proteins. Further investigation using molecularly cloned viruses revealed that a single 67D (Aspatic acid) to N (Asparagine) substitution on E protein is sufficient to confer the increased virulence and neurotropism. These findings provide new insight into Zika virus pathogenesis and suggest novel targets for the development of therapeutics.

**Author Summary:** Recent large outbreaks of Zika virus infection worldwide have revealed an association between the viral infection and increased cases of specific neurological problems including Congenital Zika Syndrome (including microcephaly) and adult Guillain–Barré Syndrome. However, the determinants of the increased neurovirulence of Zika virus remain uncertain. One hypothesis is that some unique changes across the Zika viral genome have led to the occurrence of these neurological diseases. To test this hypothesis, we continuously propagated a clinical isolate of contemporary Zika virus (SW01) in neonatal mice brain for 11 times to obtain an mouse central nervous system (CNS) adapted Zika virus (MA-SW01) that showed significantly increased neurovirulence *in vivo*. We then discovered that a single G to A nucleotide substitution at the 1069 site of Zika virus open reading frame leading to a D (aspartic acid) to N (asparagine) in viral Envelope protein is responsible for the increased neurovirulence. These findings improve our understanding of the neurological pathogenesis of Zika virus and provide clues for the development of antiviral strategy.

## Introduction

Zika virus (ZIKV) is a reemerging arborvirus that has gained worldwide attention since its large outbreaks in Southern America and rapid spread to other continents during 2015-1016. The virus was first isolated in the Zika forest of Uganda in 1947, caused only a handful of documented self-liming mild febril illness during a period of sixty years, and thus had long been neglected as a human pathogen until causing major epidemics in Yap Island in 2007 (1), French Polynesia in 2013 (2, 3), and Americas since 2014 (4). These recent large epidemics revealed some previously unappreciated facts that Zika virus infection is strongly associated with increased incidence of microcephaly in newborns, Guillain–Barré syndrome in adults, and persistent infection in male genital tissues and organs (4, 5). Among these severe complications, the link between congenital Zika virus infection and birth defect or neurologic disorder in infants has made the strongest psychological impact on the public (6–11).

The long quiescent period followed by sudden major Zika outbreaks has been postulated to be the results of viral sequence mutaions, increased competence of mosquito vector, and widely available susceptible populations (12). The interaction between Zika virus and its host has been an area of intensive study, in which specific links have been revealed, some of them emphasize the influences of viral genetics. An evolutionary Alanine to Valine (A188V) mutation on the NS1 protein of Zika virus increases virus infectivity in mosquitoes and also helps to evade interferon induction in murine cells *in vitro* and mouse model of infection *in vivo* (13, 14). A Serine to Asparagine (S139N) mutation on the prM protein of Zika virus strains isolated between 2013-2014 has been associated with neurovirulence (15). By comparing functional differences between African and Asian Zika virus strains, however, it was found that the Asian strains responsible for the recent outbreaks and linked to microcephaly do not have more infectivity to neuronal cells in cell culture *in vitro* (16), or more neurovirulence *in vivo* than the older African strains (17, 18). Other studies have placed more emphasis on the global interations between virus and host. It has been reported that high levels of virus RNA can persist in human fetal and neonatal central nervous system (CNS) *in vivo* (19, 20), and experimentally infected fetal neurocytes *in vitro* (21). Preexisting anti-flavivirus immunity can worsen clinical outcomes, through antibody dependent enhancement (ADE) of Zika virus in murine models (22–24), but not in non-human primate models (25, 26). Overall, the existing literature concurs that Zika virus infection can cause disorders in fetal and neonatal central nervous system, but reveals uncertainty on the specific virus genetic features resulting such disease outcomes.

To study these questions on Zika virus pathogenesis, various animal models have been developed, most of which use immune deficient mice that are susceptible to Zika virus infection *in vivo* such as A129 (129 background, deficient in IFN-α/β receptor), AG129 (129 background, deficient in IFN-α/β and IFN-γ receptors), A6 (C57/BL6 background, deficient in IFN-α/β receptor), AG6 (C57/BL6 background, deficient in IFN-α/β and IFN-γ receptors); Stat2 deficient mice, and Rag1 deficient mice (13,27–30). Zika virus can also infect wild type neonatal mice and cause diseases that resemble to some extent microcephaly, paralysis, and seizure (31–33). Mechanistically, these neurological manifestations have been linked to Zika virus infection of neuron progenitor cells and other neurocytes in neonatal mice *in vivo* (11,17,34). Of note, comparative analysis between human and rodents has shown that mouse brain at postnatal day 1-2 roughly corresponds to the human fetal brain at mid-gestation stage (35, 36). Therefore, newborn mice have also been used as a model for studying the influence of Zika virus infection on CNS development and pathogenesis.

Combining animal study and viral genetic analyses, here we report the isolation and characterization of Zika virus variants accumulated during sequential *in vivo* passage of a clinical isolate of Zika virus (SZ-WIV01) in neonatal mice brain. Significantly, a viral variant with a single nonsynonymous nucleotide mutation on position 1069 of Zika virus open reading frame (ORF) (G1069A), causing an amino acid mutation (D67N) on the E protein, is sufficient to account for 100-1,000 fold increase in neurovirulence in neonatal mice. These data provide increased understanding of Zika pathogenesis.

## Results

### *In vivo* adaptation of a clinical Zika virus isolate SW01 in neonatal mice

Zika virus SW01 (SZ-WIV01 strain) is a clinical isolate recovered in 2016 from a Chinese patient returned from an epidemic region, Samoa (37), expanded on C6/36 cells, titrated on Vero cells, and stored at -800C until use. To generate a mouse-adaptive viral strain, we serially passaged Zika virus SW01 in neonatal mice (**Fig. 1A**). Two day post-neonatal (DP2) mice were injected with 1,000 PFU of virus by intracranial (i.c.) route. Brains of infected mice were collected from days 2 to 12 post infection, minced, filtered, and then used for measuring virus titer with standard plaque assay. *In vivo* virus growth kinetics results showed that Zika virus SW01 replicates to a peak level of 10^6^ PFU/ml at day 8, and then gradualy decline to around 10^4^ PFU/ml at day 12 (**Fig. 1B**). And thus the day 8 viral stock was used to infect new DP2 neonatal mice from which viruses were harvested from mouse brain on day 8 and used to perform the next round of infection. After repeating this process for 11 rounds, we obtained a mouse adaptive virus of the SW01 strain, herein named MA-SW01(or MA-P11) (**Fig. 1C**).

**Figure 1.**
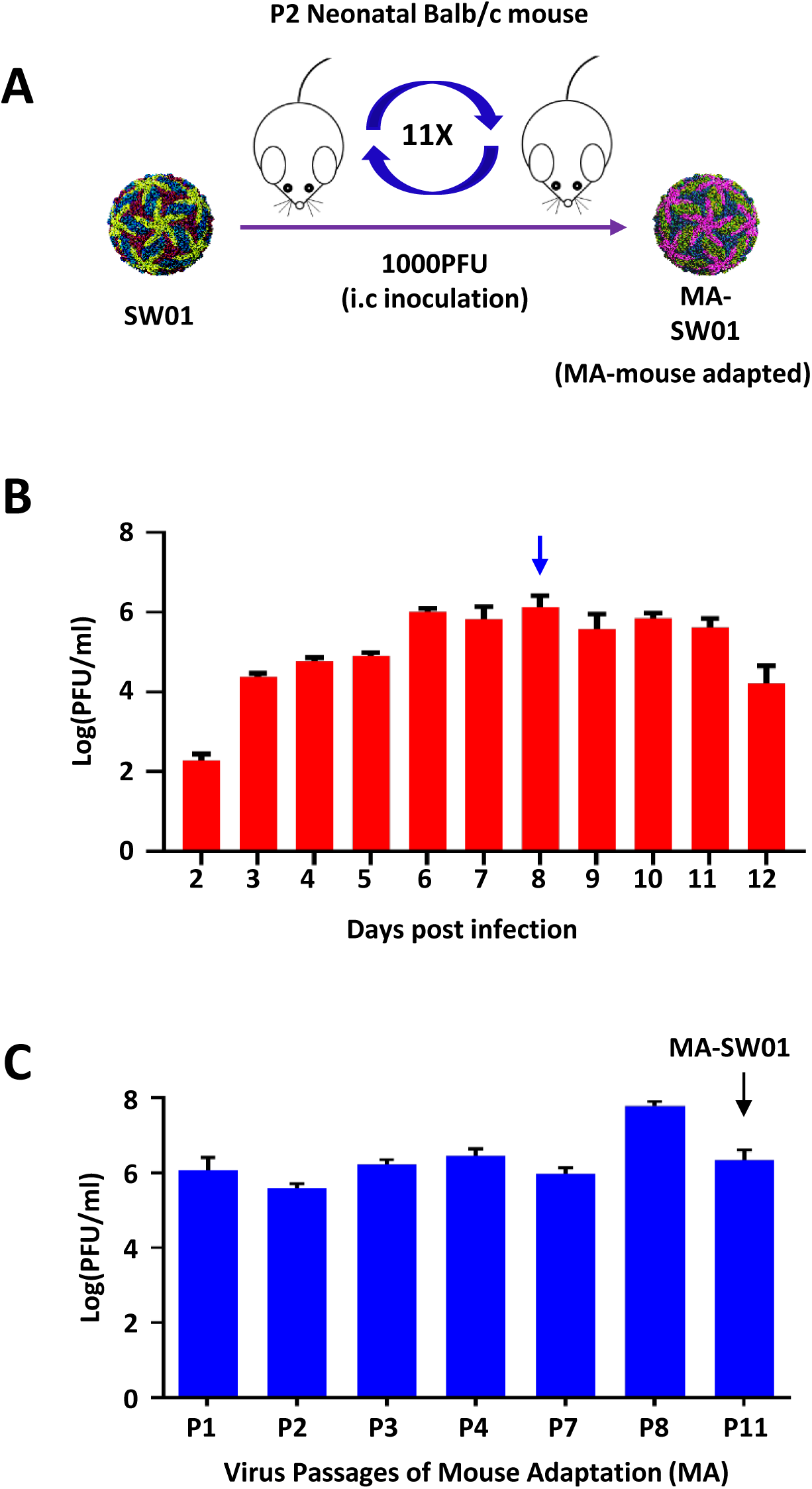
*In vivo* adaptation of Zika virus clinical isolate SW01 in neonatal mice. **(A)**. DP2 (2 days postnatal) Balb/c mice were intracranially (i.c.) infected with 1,000 PFU SW01. Brains were collected from day 2 to day 12 after infection and homogenized. Virus titers of brains were tested by standard plaque assay. **(B).** Schema of Zika virus *in vivo* passaging model; Homogenate supernatant of infected mouse brain at 8 dpi was collected and used for the next round infection in naïve DP2 Balb/c mice. This process was repeated for 11 rounds to obtain a mouse adaptive virus MA-SW01. **(C).** Virus titers from MA-P1 to MA-P11 (MA-SW01) were determinated by standard plaque assay. The summary data were presented as mean ± standard deviation (SD).

### MA-SW01 virus is more virulent than its parental virus in neonatal mice

To characterize the mouse adapted Zika virus, 100 PFU of parental SW01 virus, mouse adapted MA-SW01 virus, or negative control sterile PBS were i.c. injected into newborn DP2 Balb/c mice and monitored for up to 25 days. All mice in the SW01 group showed a slow and moderate disease progression within 11-25 days (**Fig. 2A, 2C, 2D**), in comparison, those in the MA-SW01 group showed more rapid weight loss, severe morbidity, and even death within 6-8 days (**Fig. 2B, 2C, 2D**); indicating MA-SW01 is more virulent than SW01. To determine whether the above effects are viral specific, a dose-response experiment was performed. Different groups of DP2 Balb/c mice were i.c. inoculated with 0.1, 1, or 10 PFU of SW01 virus or mouse adapted MA-SW01 virus, or sterile PBS control, and then monitored for 25 days. Results showed that at 1 or 10 PFU of MA-SW01, 100% of mice died within 7-9 days; in contrast, 1 PFU of SW01 was not lethal, and 10 PFU of SW01 only caused 77.8% fatality within a much longer period of 22-23 days. Even at 0.1 PFU, 62.5% mice in the MA-SW01 group died within 10-11 days post infection, and the remaining 37.5% survived for at least 25 days, which was the entire duration of observation; the same dose of 0.1 PFU of SW01 did not cause any death (**Fig.S1A**). These data demonstrated that MA-SW01 impacts on pathogenesis in a dose dependent manner.

**Figure 2.**
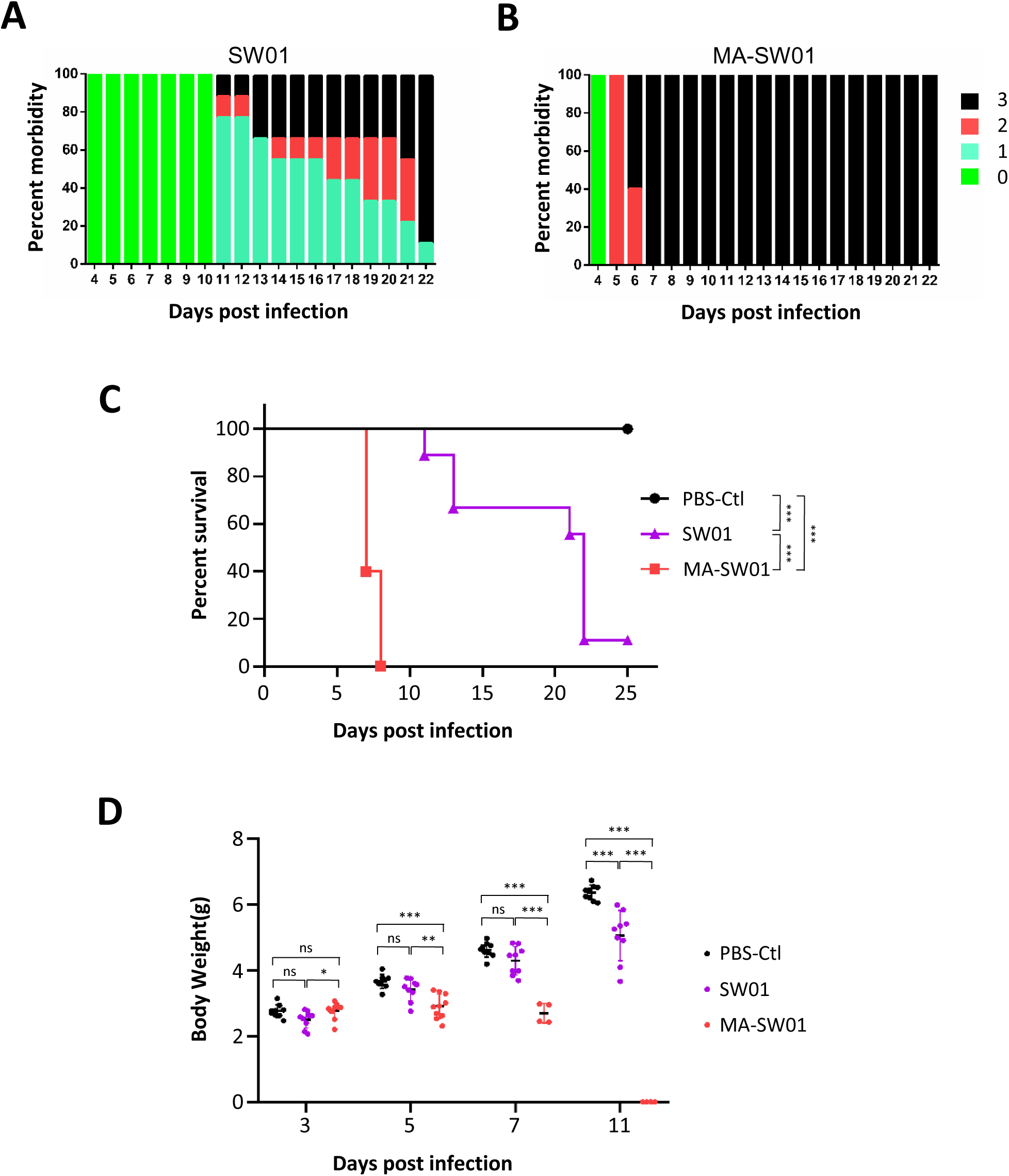
Adapted MA-SW01 virus is more virulent than its parental virus SW01. **(A-D)**. DP2 Balb/c mice were injected i.c. with 100 PFU SW01, MA-SW01, or PBS. **(A-B).**The morbidity of SW01 and MA-SW01 infected mice (Clinical Score: 0-health, 1-Manic and limb weakness, 2-limb paralysis, 3-Moribund or death);**(C).** Survival was monitored from 0 to 25 days post infection; **(D).** Body weight was analyzed at 3, 5, 7, and 11 days post infection. The summary data were presented as mean ± standard deviation (SD). Survival rate and body weight were analyzed by log rank test and two-way ANOVA respectively; P values were indicated by * (p<0.05), or ** (p<0.01), or *** (p<0.001).

To mimic natural Zika virus infection, 100 PFU of the parental SW01 virus or mouse adapted MA-SW01 virus was injected subcutaneously (s.c.) to DP2 and DP7 Balb/c mice and monitored for survival and body weight. Results showed that infection by SW01 virus was nonlethal to either DP2 or DP7 mice; in contrast, inoculation with the MA-SW01 virus caused 100% mortality in DP2 mice within 6-8 days, and 25% death in DP7 mice at 15 days post infection (**Fig.S1B**). Toghther with previous results (**Fig. 2C**), these data indicate that the virulence of MA-SW01 is age-dependent, but not related to the route of infection. To examine whether the increased virulence of the mouse adapted MA-SW01 virus is limited to only one specific mouse strain, we next injected s.c. to DP2 C57/BL6 mice with either SW01 or MA-SW01 virus, or sterile PBS as control, and then monitored them for 15 days. Results showed that MA-SW01 infected C57/BL6 mice exhibited 100% mortality at 6-7 days post infection, whereas only 44.4% of SW01 infected mice died at 15 days post infection (**Fig.S2**). Thus, the increased virulence of mouse adapted MA-SW01 virus is not restricted to one specific mouse strain.

### Increase virulence of MA-SW01 is associated with greater viral replication in multiple organs including brain

Previous studies have shown that a low-dose of Zika virus infection (2 x 10^3^ PFU) in C57/BL6 neonates led to a limited but detectable level of infection in mouse brain, and a lower mortality rate than infection of immune deficient A6 mice (IFNα/βR^-/-^, C57/BL6 background) (31). In comparison, a high-dose of Zika virus infection (10^6^ TCID50) in P1 neonatal C57/BL6 mice caused systemic infection and 100% death (32). To examine whether the MA-SW01 virus has adapted to replicate more efficiently in mice to cause systemic infection, and consequently increased virulence, DP2 Balb/c mice were infected s.c. with 100 PFU of SW01 virus or MA-SW01 virus, and then the viral loads in tissues including brain, eyes, blood, spleen, and kidney were quantified by real time-qPCR at 3 and 6 days post infection. Results showed that higher level of Zika virus RNA was detected in the brain of MA-SW01 infected mice at 3 days post infection, and higher viral loads in multiple tissues (brain, eye, blood and spleen) were observed in MA-SW01 infected mice than SW01 infected mice at 6 days post infection (**Fig. 3A**). Specifically, the average viral RNA level in the brain of MA-SW01 infected mice was about 15-fold and 488-fold higher than that of SW01 infected mice at 3 and 6 days post infection, respectively; in eyes, the average viral RNA level of MA-SW01 infected mice was 22-fold higher than that of SW01 infected mice at 6 days post infection; in spleen, the average viral RNA level of MA-SW01 infected mice exhibited approximately 5-fold increase compared to SW01 infected mice at 6 days after infection; in blood, the average viral RNA level of MA-SW01 infected mice was 54-fold higher than mice infected by SW01 virus at 6 days post infection(**Fig. 3A**).

**Figure 3.**
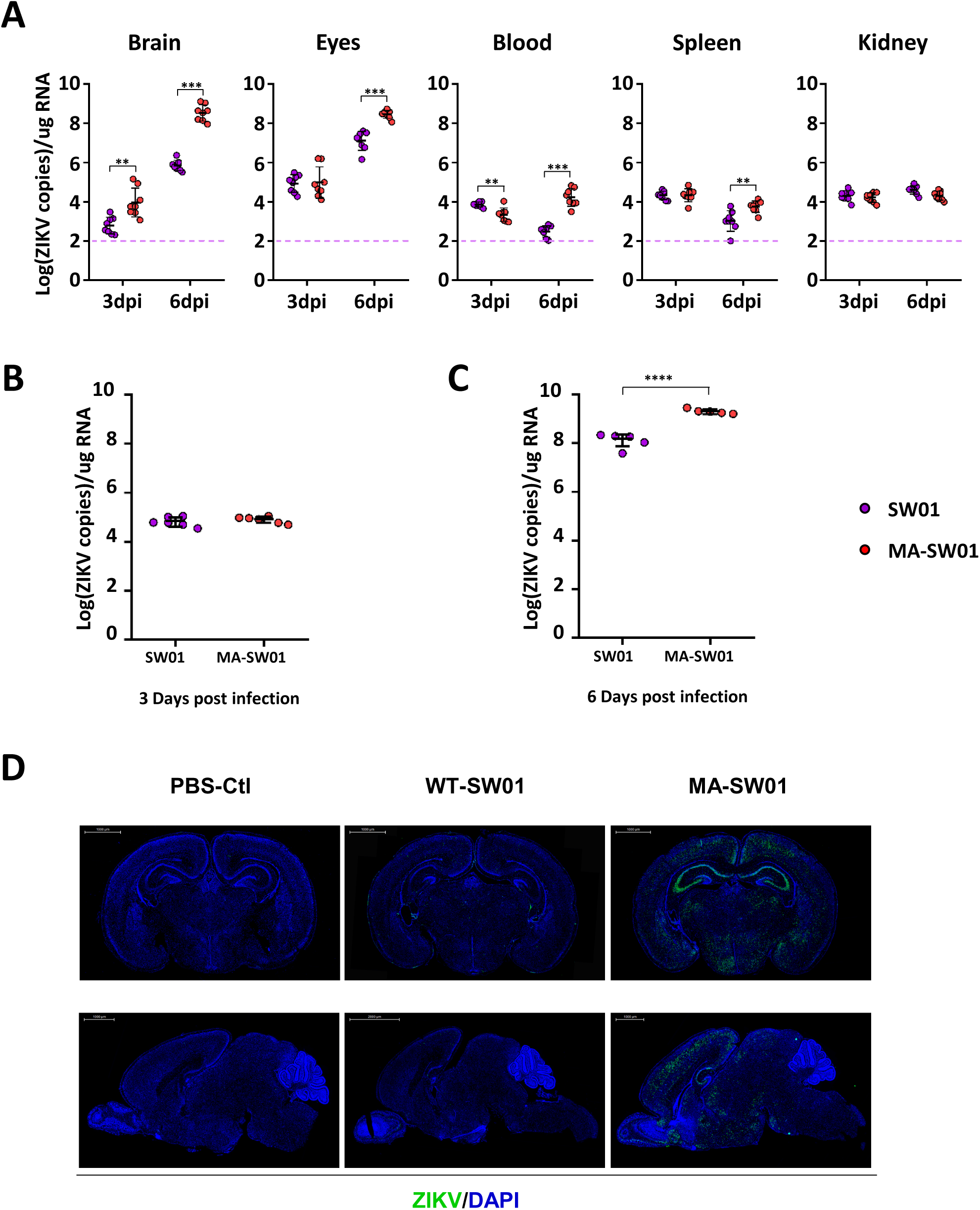
Increased virulence of MA-SW01 is associated with elevated neurotropism. **(A)**. DP2 Balb/c mice were infected s.c. with 100 PFU SW01 or MA-SW01 virus. Virus RNA loads in tissues (brain, blood, spleen, liver and kidney) at 3 days post infection (3 dpi) and 6 days post infection (6 dpi) after infection were determinated by real-time PCR; dotted lines denote the limit of detection of the real-time PCR**. (B-C).** DP2 Balb/c mice were infected i.c. with 100 PFU SW01 or MA-SW01 virus. Virus loads of brain at 3 dpi and 6 dpi was determined by real-time PCR; **(D).** DP2 Balb/c mice were infected s.c. with 100 PFU SW01 or MA-SW01 virus. Virus E protein expression in whole brain (both coronal and sagittal dissection) at 6 days post infection was detected by fluorescence immunoassay (IFA); The summary data were presented as mean ± standard deviation (SD) and analyzed by student’s t test; P values were indicated by * (p<0.05), or ** (p<0.01), or *** (p<0.001).

It is interesting to note that the average viral copy number in the brain of MA-SW01 infected mice was dramatically higher (about 488-fold) than that of SW01 infected mice at day 6 post infection (**Fig. 3A**). More viruses in the brain may be explained by two possibilities, one is that MA-SW01 virus replicates more efficiently than SW01 in the central nerve system (CNS); another is that MA-SW01 virus has increased neuro-invasion efficiency. To investigate these two possibilities, DP2 Balb/c mice were infected with 100 PFU SW01 virus or MA-SW01 virus by intracranial (i.c.) inoculation and viral RNA in the brains were quantified by real-time qPCR. Results showed that viral RNA of MA-SW01 group was not significantly different from that of SW01 group at 3 days post infection (**Fig. 3B**), but 13.8 fold higher than that of SW01 group at 6 days post infection (**Fig. 3C**), suggesting more efficient replication of MA-SW01 in the brain. Given that viral RNA level in the brain of MA-SW01 group was about 15 fold higher than that of SW01 group at day 3, even with s.c. inoculation (**Fig. 3A**), we deduced that the more virulent MA-SW01 virus has great penetration to brain. To more directly visualize viral infection in the brain, immunofluorescence staining of virus E protein in brain tissue sections was performed. Dramatically stronger fluorescent intensity indicating Zika virus E protein was detected in MA-SW01 infected mouse brains at 6 days post infection, compared to that inSW01 infected mice (**Fig. 3D**). More detailed brain staining analyses showed that cortex and hippocampus regions are major sites for MA-SW01 infection (**Fig.S3A, S3B, and S3C**), albeit the specific target cells in these tissues are currently unclear. Collectively, the above data indicate that MA-SW01 virus also has increased tropism to neuronal tissues.

### The MA-SW01 virus has four high frequency nonsynonymous mutations

To exploit whether the increased virulence of mouse adapted MA-SW01 virus is the result of unique genetic characteristics or quasispecies properties, we compared the MA-SW01 virus with its parental virus SW01 at the phenotypic and genetic levels. Biological clones derived from MA-SW01 virus (MA-1, MA-2, MA-3, MA-4, MA-5, MA-6, MA-7, MA-8, MA-9 and MA-10) were found to be more virulent than clones from SW01 (SW-1, SW-2, SW-3, and SW-4) (**Fig.S4**), indicating the increased virulence of MA-SW01 is not a reflection of viral quasispecies but may be related to specific genetic characteristics. To uncover the genetic changes during *in vivo* adaptation that might have caused the increased virus virulence, RNA was extracted from the original SW01 viral stock and mouse adapted MA-SW01 virus, and then subjected to sequencing by the next generation sequencing (NGS) method. Intrahost single nucleotide variant (iSNV) across the whole genome was analyzed by CLC genomic workbench.

Results showed 6 nucleotide substitutions (G1069A, G1074A, C1089T, A1330G, G3787A and T6036C) over 80% reads in the ORF (open reading frame) of mouse adapted MA-SW01 virus (**Fig. 4A**). Among these substitutions, four were nonsynonymous mutations (G1069A, G1074A, A1330G and G3787A), and two were synonymous mutations (C1089T and T6036C); the four nonsynonymous mutations led to three amino acid changes on virus E protein (D67N, M68I, N154D), and one on NS2A protein (A117T) (**Fig. 4B**). Of note, N154 is a unique glycosylation site on Zika virus E protein, and it has been shown to support Zika virus infection in adult immune-deficient mice by either enhancing virus neuroinvasion or facilitating DC-SIGN binding (38, 39). However, when Zika virus was inoculated intracranially in neonatal mice, the deletion of N154 glycosylation had no impact on virus virulence (40). Collectively, these published data suggest that the N154D mutation may not be linked to the augmented infectivity and increased neurovirulce we have observed for the mouse adapted MA-SW01 virus. Hence, we focused on the other 3 amino acid mutations (D67N, M68I on E protein, and A117T on NS2A protein) for further study.

**Figure 4.**
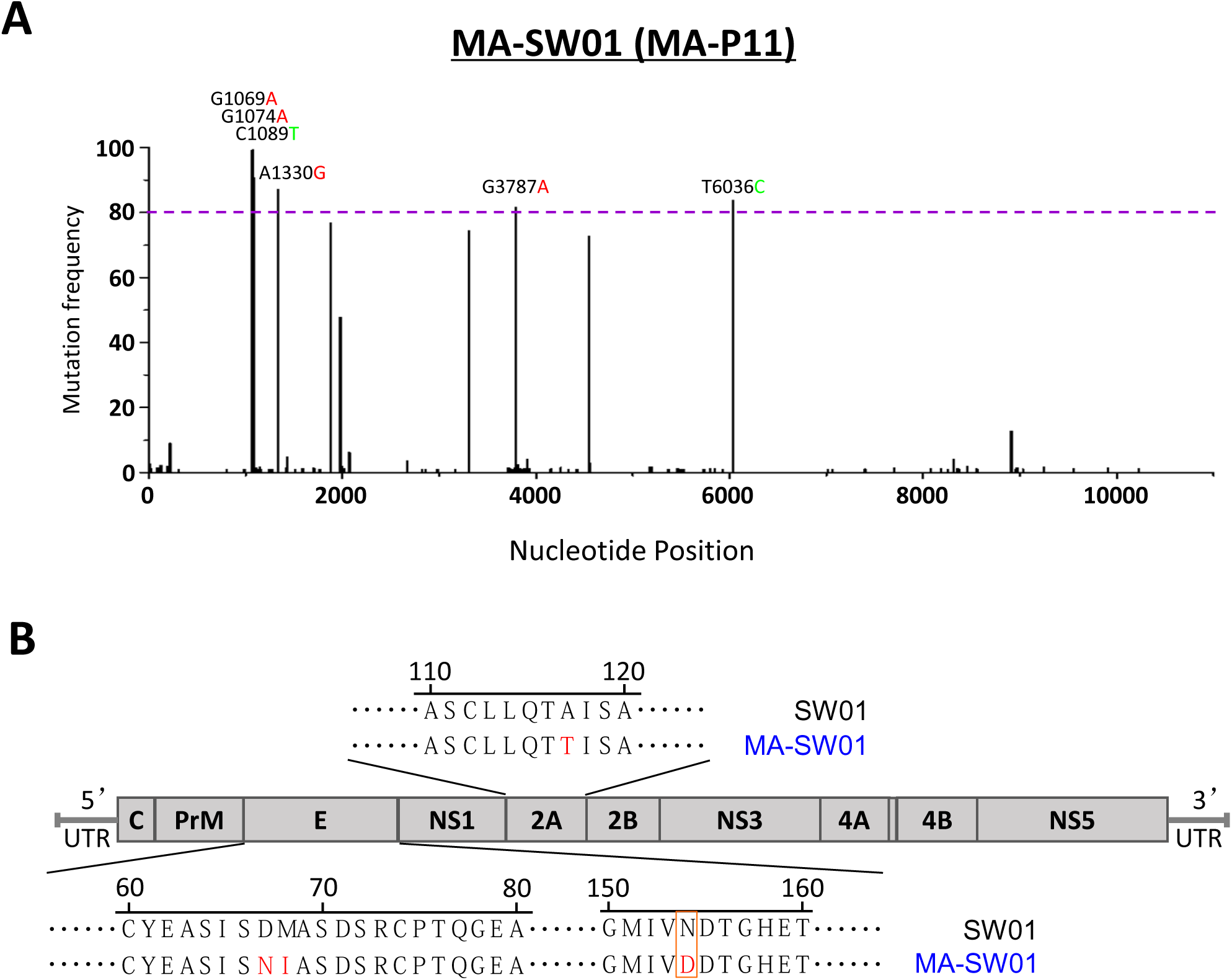
NGS analyses of the MA-SW01 virus identify 4 high frequency nonsynonymous mutations on E and NS2A genes. **(A-B)**. Virus RNA extracted from SW01 and MA-SW01 was used to construct the sequence library, and then sequenced by the next generation sequencing (NGS) method. Quantification and plotting of mutation frequency were performed by CLC Genomic Workbench and Origin software. **(A).** Plots of missense mutations frequency across the ORF of MA-SW01 in reference to consensus sequence of SW01; Nucleotides with frequency higher than 80% reads were shown (Red nucleotide abbreviations represent missense mutations, and green ones are silent mutations). Dotted lines denote the frequency of 80%. **(B).** Amino acid changes corresponding to missense mutations.

### Mutations on E protein are required for increased virulence

Based on a widely used Zika virus infectious clone pFLZIKV (41), three mutant viruses (CM1, CM2 and CM3) were constructed: CM1 includes D67N and M68I mutations on E protein; CM2 includes the A117T mutation on NS2A protein; CM3 contains all three substitutions on E and NS2A proteins (**Fig. 5A**). In BHK-21 cells, 1-3 days after transfection with *in vitro* transcripted RNA from these viral constructs, virus E protein expression was examined with a monoclonal antibody. Results showed that all three molecularly cloned mutate viruses were rescued and replicated efficiently (**Fig. 5B**). These molecularly cloned mutant viruses (CM1, CM2, and CM3 virus) were then used to infect DP2 Balb/c mice i.c. at 10 PFU/mouse, using the parental molecularly cloned CAM-WT virus as a control, and then monitored for 25 days. Results showed that CM1 and CM3, but not CM2, were more virulent than parental virus CAM-WT in neonatal mice (**Fig. 5C**). Since both CM1 and CM3 contain 2 mutations on E protein (D67N, M68I), and CM2 only contains the single NS2A mutation, the above results suggest that the E protein mutations were determinants of increased virulence of MA-SW01, and the NS2A mutation was not. Therefore, CM1 virus containing two E protein mutations was chosen for further investigation.

**Figure 5.**
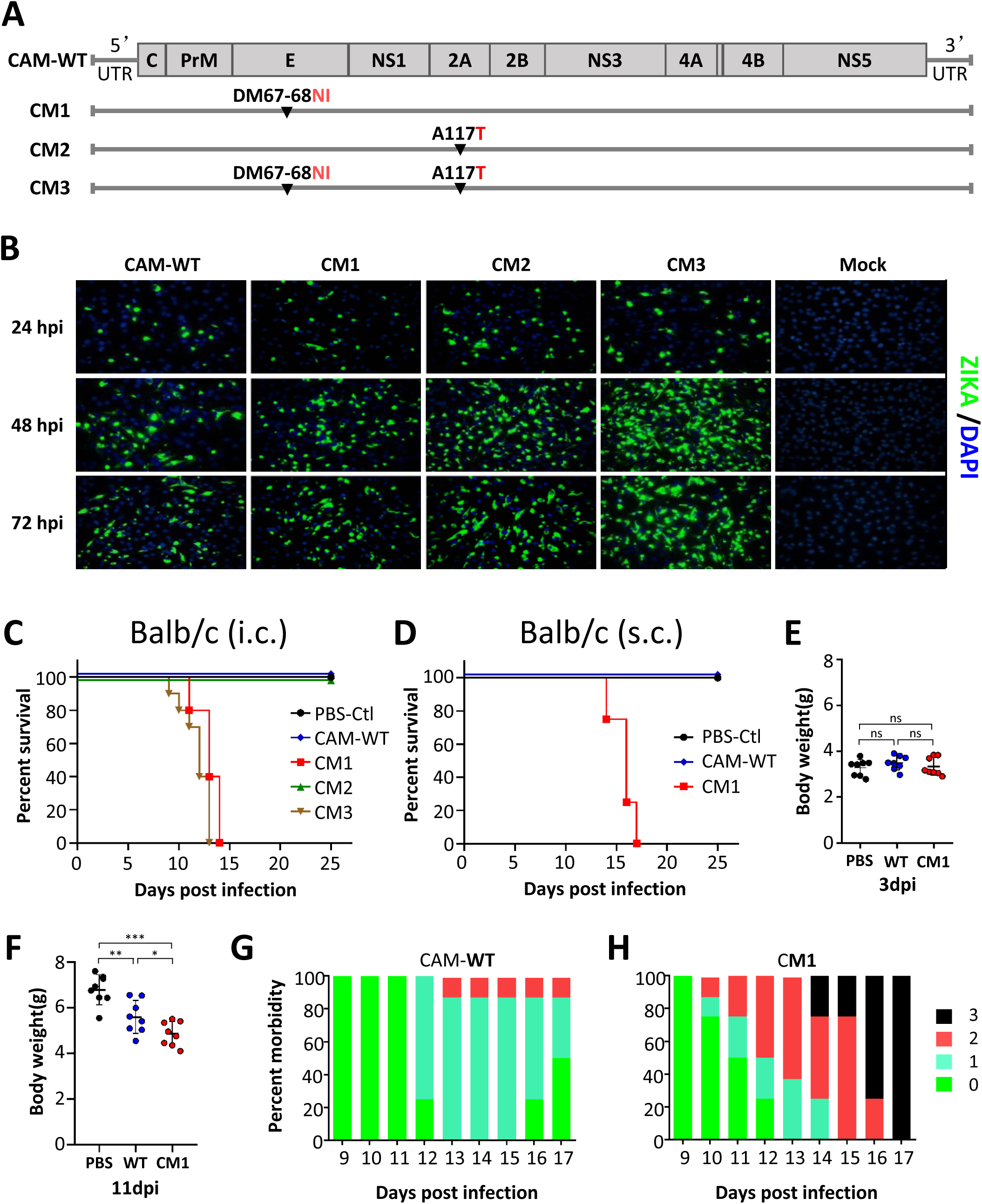
Increased virulence of Zika virus is associated with specific mutations on E protein but not on NS2A protein. **(A)**. Scheme of mutation strategy based on pFLZIKV (CAM-WT) infectious clone to create CAM-M1 (CM1), CAM-M2 (CM2), CAM-M3 (CM3) viruses; **(B).** IFA of Zika virus E protein expression at indicated times (24, 48, 72 hours post infection) in BHK-21 cells transfected with RNA from CAM-WT or mutant viruses (CM1, CM2, CM3). **(C).** Survival curve of DP2 Balb/c mice infected i.c. with 10 PFU CAM-WT, mutant viruses (CM1, CM2, CM3), or PBS. **(D-H).** DP2 Balb/c mice were infected s.c. with 100 PFU CAM-WT, mutant viruses, or PBS. **(D).** Survival was monitored and analyzed from 0 to 25 days post infection; **(E**-**F).** Body weight difference between PBS-Ctl (PBS), CAM-WT (WT) and CAM-M1 (CM1) groups at 3 and 11 days post infection (3 dpi and 11 dpi); **(G-H).** The morbidity of CAM-WT and CAM-M1 groups (clinical score: 0-health, 1-manic and limb weakness, 2-limb paralysis, 3-moribund or death); the summary data were presented as mean ± standard deviation (SD) and analyzed by student’s t test; P values were indicated by * (p<0.05), or ** (p<0.01), or *** (p<0.001).

We first examined whether the route of infection alters viral virulence. DP2 Balb/c mice were infected s.c. with 100 PFU of control CAM-WT or test CM1 virus, and then monitored for up to 25 days. Results showed that CM1 infection led to higher motality than that of CAM-WT (**Fig. 5D**). Although a difference was not observed earlier at 3 days post infection (**Fig. 5E**), body weight recorded at 11 days post infection showed that CM1 infected mice were significantly lighter than those infected by CAM-WT (**Fig. 5F**). Notably, disease progression appeared to be reversible in the control CAM-WT group, but not so in the CM1 group which had 100% mortality at 17 days post infection (**Fig. 5G, 5H**). These data demonstrated that the severe outcome as a result of CM1 infection is not constrained by inoculation routes.

Since virulence of the mouse adapted MA-SW01 virus was not restricted to a single mouse strain, we sought to confirm that the molecularly cloned CM1 viurs follows the same principle. DP2 C57/BL6 mice were inoculated s.c. with 100 PFU of parental CAM-WT or test CM1, or negative control PBS, and then monitored for 25 days. All mice (100%) infected by CM1 virus died at 11-13 days post infection, whereas only 20% of those infected by CAM-WT succumbed at 25 days post infection (**Fig.S5**). These data demonstrated that the virulent phenotype of CM1 is not mouse strain specific.

### A D67N single mutation is sufficient to account for the increased virulence of the molecularly cloned CM1 virus

To determine which one of the two amino acids is more critical for a major change in viral phenotype, we first analyzed single nucleotide variants in E protein sequence from serially passaged P1 (MA-P1) to P11 (MA-P11) viruses. Results revealed that there were progressive accumulations of D67N (from 22.7% to 99.4%), and M68I (from 3.0 % to 91.7%) mutations during the *in vivo* serial passage of parental SW01 virus. The baseline frequencies of these two mutations in the parental virus (P0, or SW01) were lower than 1%. Of note, the D67N maintained high mutation frequency (>90%) from P5 to P11 during *in vivo* passage (**Fig. 6A**). Consistent with the notion that this mutation may be functionally significant, DP2 C57/BL6 mice infected with 100 PFU of MA-P5, MA-P8 or MA-P10 showed 100% mortality at 10 days post infection, whereas the parental virus SW01 infected mice had only 16.7% mortality at 15 days post infection (**Fig. 6B**). Given that the D67N mutation rapidly increased to 92.4% in P5 virus (**Fig. 6B**), and all single biological clones of the mouse adapted MA-SW01 virus contain the D67N mutation (**Table.S1**), it is reasonalb to deduce that D67N mutation alone is responsible for the increased viral virulence. To test this idea, a molecular clone contains the single D67N mutation was constructed based on the CAM-WT backbone (pFLZIKV), and herein named CM1-A virus (**Fig. 6C**), which was rescued successfully as showed by immunofluorescence staining of ZIKA VIRUS E protein of BHK-21 cells transfected with *in vitro* transcripted RNA from CAM-WT or CM1-A viruses (**Fig. 6D**). Then, 100 PFU of CM1, CM1-A virus or PBS was inoculated s.c. into DP2 C57/BL6 mice which were then monitored for up to 25 days. Results showed that CM1-A was similar to CM1 in causing 100% mortality of infected mice at 12-13 days post infection (**Fig. 6E**). These data demonstrated that a single D67N mutation is sufficient to account for the increased virulence of CM1.

**Figure 6.**
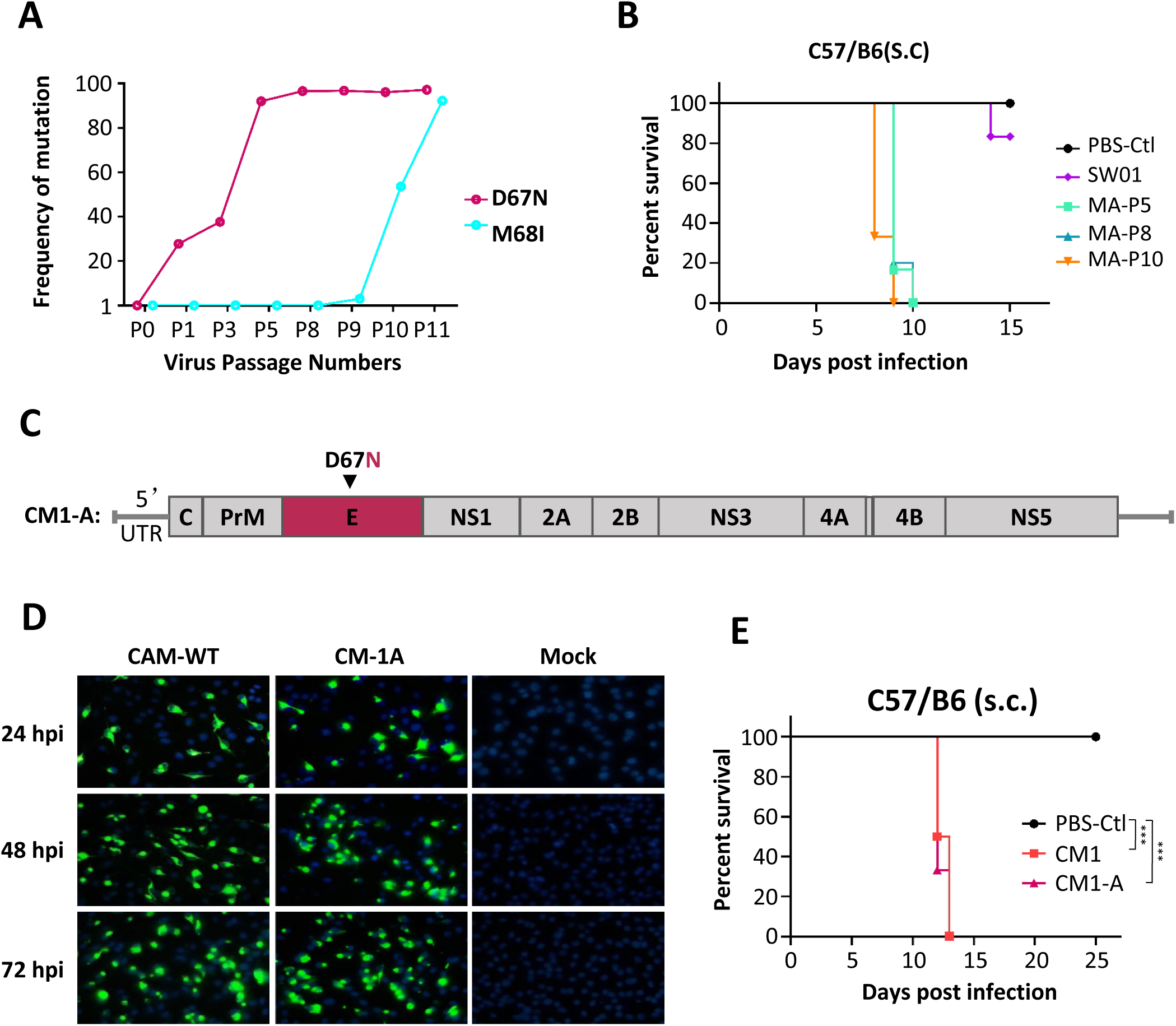
Increased virulence of Zika virus is associated with a single D67N mutation on E protein. **(A)**. Progressive changes of mutation frequency of 67 and 68 amino acids in E protein from SW01 (P0) to MA-SW01 (MA-P11) during *in vivo* passaging. **(B).** Survival curve of DP2 C57/BL6 mice infected s.c. with 100 PFU SW01, MA-P5, MA-P8, MA-P10, or PBS. **(C).** Strategy to construct a single D67N substitution virus (CM1-A) based on CAM-WT infectious clone; **(D).** IFA of Zika virus E protein expression at indicated times (24, 48, 72 hours post infection) in BHK-21 cells transfected with RNA from CAM-WT and CM1-A. **(E).** Survival curve of DP2 C57/BL6 mice inoculated s.c. with 100 PFU CM1, CM1A virus, or PBS. Survival rate was analyzed by log rank test; P values were indicated by * (p<0.05), or ** (p<0.01), or *** (p<0.001).

### D67N mutation on E protein promotes Zika virus infection in brain

To investigate whether viral E protein mutations (D67N, M68I) influence tissue tropisms, DP2 Balb/c mice were infected s.c. with either parental CAM-WT or test CM1 (containg both D67N, M68I), then euthanatized at 3 and 11 days post infection. Viral RNA in tissues were quantified by standard real-time qPCR. At 3 days post infection, all tissues except for eyes showed similar levels of viral RNA between CAM-WT and CM1 groups; at 11 days post infection, however, viral RNA of CM1 group was significantly higher than that of CAM-WT group in brains, eyes and blood, but not in spleens and kidneys (**Fig. 7A**), confirming the combination of these E protein mutations affect tissue tropism.

**Figure 7.**
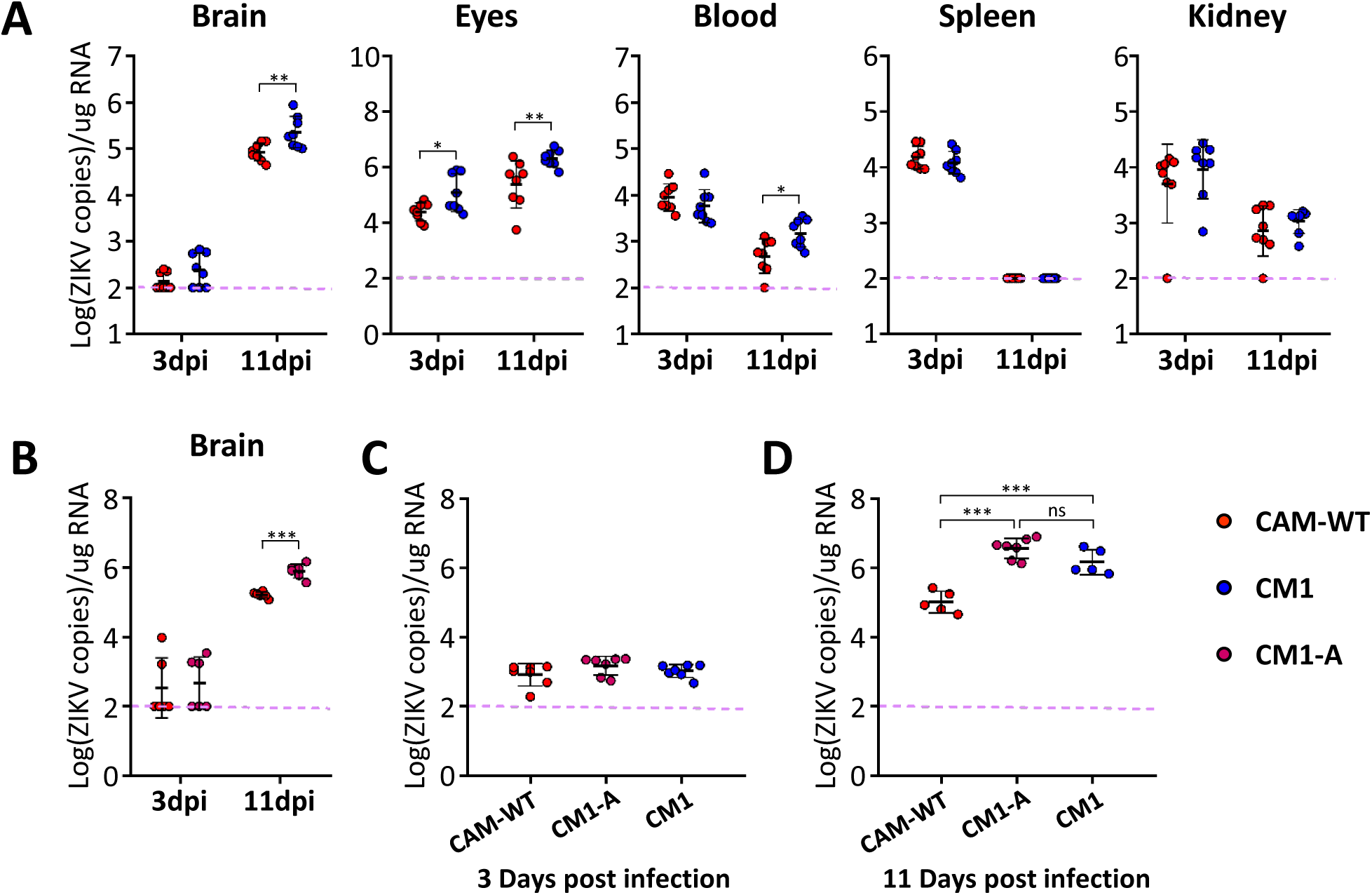
Molecularly cloned Zika virus with a single D67N mutation on E protein has marked increase in the infection of brain tissues. **(A)**. DP2 Balb/c mice were infected s.c. with 100 PFU CAM-WT or CM1. Virus RNA loads in tissues (brain, blood, spleen, liver and kidney) at 3 days post infection (3 dpi) and 11 days post infection (11 dpi) were determined by real-time PCR;**(B).** DP2 Balb/c mice were infected s.c. with 100 PFU CAM-WT or CM1-A. Virus RNA loads in brains at 3dpi and 11dpi were determined by real time PCR; **(C-D).** DP2 Balb/c mice were infected i.c. with 20 PFU CAM-WT, CM1 and CM1-A. Virus RNA load in brains at 3 dpi **(C)**and 11 dpi **(D)** were determined by real-time PCR; The summary data were presented as mean ± standard deviation (SD) and analyzed by student’s t test; P values were indicated by * (p<0.05), or ** (p<0.01), or *** (p<0.001).

The fact that more viral RNA was detected in brain and eyes, that are rich in nerve cells susceptible to Zika virus infection, implies that CM1 virus has growth advantage over CAM-WT in these cells. To directly test this notion, DP2 Balb/c mice were infected i.c. with CAM-WT or CM1, and then monitored for virus burden by real-time qPCR. Results showed that brain viral loads of CM1 group was higher than that of CAM-WT group at 11 days post infection, but not 3 days post infection (**Fig. 7C, 7D**), suggesting that CM1 has growth advantage over CAM-WT in brain.

To further dissect wthether single D67N mutation in E protein plays the essential role of altering viral virulence and tissue tropism, we next used CM1-A virus (containing only D67N) to perform viral infection experiments. Results showed that irrespective of throught s.c. infection (**Fig. 7B**), or i.c. infection (**Fig. 7C, 7D**), viral load in brains of CM1-A group were always higher than that of CAM-WT group at 11 days post infection, but not at 3 days post infection. Collectively, these data indicated that D67N mutation promotes virus virulence partially through enhancing virus replication in CNS.

### Rapid D67N accumulation may have increased virus fitness

To confirm that the observed dominant D67N substitution in the *in vivo* adapted MA-SW01 virus is originated from the parental clinical isolate SW01 virus, instead of an artifact resulted from an extraneous *in vitro* cell culture selection process, we analyzed the nucleotide polymorphism at the 1069 postion of Zika virus SW01 open reading frame (ORF), which corresponds to amino acid sequence at the 67 position of E protein. Results showed that 1069A variant of ORF was present in the initial SW01 stock at a low frequency of 0.055% (**Table.1**), indicating a minority of viral quasispecies features the N67 on its E protein. Together with previous data (**Fig. 6A, 6B**), it is reasonable to deduce that D67N substitution provides more viral fitness in CNS, and thus enables MA-SW01 to outgrow other viral variants within the SW01 quasispecies.

**Table 1.** Deduced amino acid polymorphism at the 67 site of ZIKA E protein of WT-SW01 strain

| Reads*<br>(1069 of ORF) | Nucleotide polymorphism<br>(1069 of ORF) | Amino acid polymorphism<br>(67 of Envelope) | Percentage of total Reads |
| --- | --- | --- | --- |
| 18167 | 1069 (G,T,C) | 67 (Asp, Tyr, His) | 99.945% |
| 10 | 1069 (A) | 67 (Asn)# | 0.055% |

\*: Based on deep sequencing
#: The D67N mutation presents at low frequency in the original viral isolate.

## Discussion

The pathogenesis of neurological disorders in association with Zika virus infection has been an area of intensive investigation recently. The scientific progress, however, has been hampered in part by the lack of robust *in vivo* experimental models to study the cause-and-effect between Zika virus infection and various clinical outcomes. To this end, we have generated a mouse adapted Zika virus (MA-SW01) strain that showed 100-1,000 fold increased virulence than its parental virus clinical isolate SW01, with corresponding increase in neurotropism, by serially passage of SW01 in the brains of neonatal mice. NGS analyses revealed that the MA-SW01 virus has four dominant nonsynonymous nucleotide mutations on genes encording E protein (3 mutations) and NS2A protein (1 mutation). Mechanistic studies using molecularly cloned Zika virus varients contain these muations either alone or in combination demonstate that a single nucleotide G1069A mutation in Zika virus ORF that causes an amino acid change (D67N) on the E protein is sufficient to confer greater viral replication in mouse brain and much increased mortality. These results not only establish an *in vivo* model by which many facets of Zika virus pathogenesis could be further studied, but also provide a tangible viral genetic basis to explain the neutrological complications observed in association with Zika virus infection.

Identifying viral genetical features that may affect the outcomes of viral infectios is an area of significant interest. Zika viruses with mutations on prM, NS1, NS2A have been reported to profoundly change viral infectivity in mosquitoes, cell lines, and mice (13,15,30). In the current study, we demonstrated for the first time that a single mutation on E protein can markedly increase the neurovirulence of Zika virus.

The specific mechanisms that explain why a single D67N mutation on E protein can dramatically change Zika virus infectivity is currently unknown. One reason may be its potential link to glycosylation. In all four DENV serotypes, there are two highly conserved N-linked glycosylation sites on E protein, N67 and N153, that play critical role for viral entry (42). However, similar to West Nile virus (WNV) and Japanes encephalitis virus (JEV), Zika virus has only one N154 glycosylation site on E protein (40, 43). Indeed, glycosylation on N153 or N154 of E protein helps WNV or JEV to invade CNS in mammals (38,39,44,45). Surprisingly, an additional use of N67 glycosylation on JEV attenuates virus pathogenesis and reduces viral neuro-invasion *in vivo* (45), despite the N67 glycosylation mediates enhanced infectivity of WNV or DENV by facilitating interaction with DC-SIGN molecule on cell surface in cell culture models (45–47). The influence of un-glycosylated N67 on viral pathogenesis is unknown, despite of the existence of this form of N67 on a number of flaviviruses (48). Because the mosue adapted MA-SW01 virus has a mutation on the N154 site and unable to acquire glycosylation through this site, it is tempting to suggest that the D67N is a functional compensatory mutation. It would be interesting to test whether MA-SW01 is glycosalted at the 67 site in future studies.

Other than mechanistically interesting, our findings may have practical implicaitons as well. The amino acid D67 on E protein is conserved among many known Zika virus strains, and it forms a recognition site for several monoclonal antibodies (mAbs) isolated from Zika virus infected patients; some of these mAbs can prevent Zika virus infection in animal models (49–52), indicating the 67 site being functionally important. It would be interesting to test in future studies whether our mouse adapted virus that has a D67N mutation can escape these antibodies, and thereby become more virulent. With respect to the N154 glycosylation site, our data are different from a recent study which showed that an artificial deletion of N154 glycosylation of Zika virus E protein decreases viral infectivity and neuroinvasion in mice, while maintaining viral immunogenicity *in vivo*, and thus being a promising strategy for making live attenuated vaccine (40). We found that mouse adapted Zika virus clone (MA-SW01) without the N154 glycosylation site on E protein is still highly virulent, even more so than viruses cloned from the parental SW01 virus which contains an intact N154 glycosylation site on E protein. Therefore, the vaccine stategy utilizing a deletion of N154 glycosylation may only apply to some viral variants, but not others.

In conclusion, we have identified a single amino acid D67N mutation on the putative glycosylation site of E protein of Zika virus, and demonstrated the mutant virus having profoundly increased viral virulence and neurotropism in mice. Close monitoring and large-scale screening of this unique viral variant in humans should provide clue to understand some major questions in the field, such as the sudden outbreak of Zika disease in certain locale, and the association between Zika virus infection and neurological disorders.

## Materials and methods

### Ethic statement

All experiments were performed strictly in accordance with the guidelines of care and use of laboratory animals by the Ministry of Science and Technology of the People’s Republic of China and regulations of biosafety level 2 (BSL-2) and animal biosafety level-2 (A-BSL-2) containment facilities at Institut Pasteur of Shanghai. The animal protocols were approved by the biosafety laboratory and the institutional Animal Care and Use Committee at Institut Pasteur of Shanghai (Approval number: A2018027). All mice used in this study were carefully fed and suffering of animals was minimized.

### Mouse experiments

Balb/c and C57BL/6 mice (B6) were purchased (Vital River Laboratory Animal Technology Co., Ltd, Beijing) and bred for experiments under specific pathogen free (SPF) conditions at the BSL2 Animal Core facility (A-BSL-2) at Institut Pasteur of Shanghai. Balb/c and C57BL/6 pregnant mice were housed separately before being delivered to the BSL-2 laboratory. DP2 (2 days post-delivery) and DP7 (7 days post-delivery) offspring mice were infected with indicated Zika virus strain through subcutaneous (s.c.) or intracranial (i.c.) injection. Body weight, survival rate and clinical score were monitored daily according to experimental design.

### Cell lines and viruses

Vero-E6 and BHK-21 cells were grown at 37°C in Dulbecco’s Modified Eagle Medium (DMEM) (Gibco, USA) supplemented with 10% fetal bovine serum (FBS) (Gibco, USA) and 1% penicillin and streptomycin (P/S). Mosquito C6/36 cells were cultured in Modified Eagle Medium (MEM) (Gibco, USA) with 10% FBS, 1% P/S and 1% non-essential amino acids (NEAA). Zika virus clinical isolate SW01 (also known as SZ-WIV01, GenBank: MH055376.1) was kindly provided by Wuhan Institute of Virology, Chinese Academy of Sciences. SW01 was propagated once in C6/36 cells with MEM (Gibico) plus 2% FBS, 1% Penicillin-Streptomycin and 1% non-essential amino acids. Rescued virus mutates with the backbone of Zika virus CAM-2010 infectious clone were passaged once in C6/36 cells. All amplified viruses were aliquot into 2ml vials and stocked at -80°C until use.

### Virus titration

Virus titer was determinated by titration on Vero-E6 monolayer. Briefly, Vero-E6 cells were seeded on 24 well plate (1-1.2×10^5^ cells/well) one day prior to infection, and washed once next day with DMEM without FBS. Virus was 10-fold serially diluted, then 200µl of virus was added to the Vero cell monolayer, (followed by incubation at 37°C for 2 hours.The supernatant containing virus was replaced by 1.2ml DMEM with 1.5% FBS, 1% CMC (carboxymethylcellulose), then incubated at 37°C, 5%CO2 for 96 hours. Four days later, the overlay was removed and cells were fixed with 4% PFA for 30min. The viral plaque was visualized and calculated after being stained by 0.25% crystal violet.

### Adaptation of SW01 *in vivo*

Zika virus clinical isolate SW01(10^3^ PFU/10μl) was injected into the brain λ point of Balb/c DP2 neonatal mice. At the indicated time (1-12 days and 8 days) post infection, mice were anaesthetized and brains were collected and homogenized in 1ml sterile PBS. Then, the homogenized suspension was centrifuged to collect supernatant, which was aliquot for viral titration and stored at -80°C. A new round of *in vivo* infection into the mouse brain was performed after viral titration.

### Determination of virus burden in tissues

At the indicated time (3, 6 and 11 days post infection), Zika virus infected mice were euthanized and tissues were collected, fixed with Trizol (Invitrogen, USA) reagent. RNA was extracted according to the manufactures’ manual, then aliquot and stored at -80°C before use. RNA concentration was determinated by Nanodrop 2000 (Thermo fisher, USA). Reverse transcription with ZIKA virus specific primer (Rev-AAGTGATCCATGTGATCAGTTGATCC) was performed using FastQuant RT Kit (Tiangen). Real-time PCR was done on 7900HT (ABI) machine using Fast Fire qPCR Premix (Probe) (Tiangen). Virus RNA copies were calculated with a standard curve established with NS1 gene transcript. The primers are as follows: (For: CAACCACAGC-AAGCGGAAG, Rev: AAGTGATCCATGTGATCAGTTGATCC, Probe: 5’-FAM/TGGTATGGAATGGA GATAAGGC/MGB-3’).

### Immunostaining of brain sections

Dissected brains were immediately immersed in 4% paraformaldehyde (PFA) and fixed for 24 hours. Then the fixed tissues were embedded into paraffin according to a standard protocol. Embedded brains were sectioned into 4 μm slices using Leica RM2016. After being deparaffinized with xylene and ethanol, rehydrated with ethanol and H2O, sections were blocked in blocking buffer (3% BSA-PBS) for 30 min, then incubated with primary antibody targating Zika virus envelope protein (1:1000 dilutued in blocking buffer; cat.no: BF-1176-56; BioFront)overnight at 4°C. On day 2, the sections were incubated in fluorescence labelled secondary antibody (1:400, GB25301, Servicebio) at RT for 1 hour. Nucleus were stained with DAPI (G1012, Servicebio) at RT for 10 min. Original images were captured and visualized using a Nikon Eclipse C1 Ortho-Fluorescent microscope with Nikon DS-U3 image system. All immunofluorescent images were analyzed with the Pannoramic Viewer (3DHISTECH), ImageJ, and GraphPad V8 software.

### Single clone selection and E protein sequencing

Stocks of SW01 and MA-SW01 virus were serially diluted and seeded on Vero monolayer in 24 well plate. Four days post infection, the supernatants from wells containing single virus plaque were collected and amplified in C6/36 cells once, and viral titer was determinated by standard plaque assay. For E protein sequencing, RNA of single virus clone was extracted by Viral RNA Mini Kit (QIAGEN) and reversely transcribed using PrimeScript^TM^ II 1st Strand cDNA Synthesis Kit (TaKaRa) with virus envelope protein gene specific primer (Env-Rev primer: CGGGATCCCGAGCAGAGACGGCTGTGGATAAG). Virus E gene was amplified by PCR (Env-For primer: CGAAGCTTATGATCAGGTGCATAGGAGTCAGCA, Env-Rev primer: CGGGATCCCGAGCAGAGACGGCTGTGGATAAG), then cloned into pEASY-Blunt Cloning Kit (Transgen) and sequenced by Sanger method.

### Next generation sequencing (NGS) of ZIKA virus

Briefly, virus stocks were filtered through a 0.45 μm filter before nucleic acid extraction. Virus RNA was extracted from 400 µl of filtered supernatant with the High Pure Viral RNA Kit (Roche). The sequencing library was constructed using Ion Total RNA-Seq Kit v2 (Thermo Fisher Scientific) and sequenced on an Ion S5 sequencer (Thermo Fisher Scientific). Low quality reads and short reads were filtered. All filtered reads were assembled by mapping to the reference sequence MH055376 using CLC Genomic Workbench (ver 9.0). The mutation site was manually checked with original sequencing data. iSNV and Graphing were performed on CLCGenomic Workbench and Origin. NGS raw data were available at Sequence Read Archive (SRA) of NCBI (Access number: SRP237251).

### Generation of Zika virus mutants

The infectious cDNA clone (pFLZIKV) containing Zika virus CAM-2010 full-length genome were used as the backbone for introducing the nucleotides substitutions into envelope protein (D67N, M68I) or NS2A protein (A117T), singly or combined, using the Q5 site directed mutagenesis kit (NEB). All the mutations were confirmed by DNA sequencing. The full-length infectious clones were rescued as described previously (41).

### Indirect immunofluorescence assay (IFA)

The viral RNA was transfected into BHK-21 cells using Lipofectamine 3000 reagent (Thermo Fisher Scientific) according to the manufacturer’s instructions. At 24, 48, and 72 hr post infection, the infected cells were fixed in acetone/methanol (V/V=3/7) at -20 °C for 15 min, and then used for detection of Zika virus E protein expression by IFA as described previously (53).

### Statistic analysis

Survival curves were analyzed by log rank test. Body weight was analyzed by Two-way ANOVA (Turkey correction). All summarized data were compared for statistical differences by student’s t test or two-way-ANOVA. All analyses were performed on Graphpad Prism V8.0 platform. Statistical significance levels were reported as the following: “*” for p < 0.05; “**” for p < 0.01; “***” for p < 0.001 or less.

## Acknowledgement

We thank Dr. C.Y Zhang (Institut Pasteur of Shanghai, Shanghai, China) for kindly providing information on primers and RNA standard used for Real-time PCR assay of ZIKA virus. The study was supported in part by the following grants: Strategic Priority Research Program of the Chinese Academy of Sciences (XDB29040301, X.J.), National Key R & D Program of China (2016YFC1201000, X.J.), Ministry of Science and Technology of China (2016YFE0133500, X.J.), European Union Horizon 2020 Research and Innovation Programme under ZIKAlliance Grant Agreement 734548 (X.J.).

## Author Contributions

X.J. supervised the research; X.J., ZH.L. conceived the research; ZH.L., YW.Z., ML.C., YG.T., CF.Q., X.J. contributed to the project design and results discussion; ZH.L., YW.Z., ML.C., NN.G. and JY.S. performed the experiments and analyzed the experimental data; ZH.L. and YW.Z. analyzed and summarized the NGS data. ZH.L. wrote the original manuscript. ZH.L., YW.Z., ML.C., YG.T., CF.Q. and X.J. revised and edited the manuscript.

